# Eukaryotically expressed encapsulins as orthogonal compartments for multiscale molecular imaging

**DOI:** 10.1101/222083

**Authors:** Felix Sigmund, Christoph Massner, Philipp Erdmann, Anja Stelzl, Hannes Rolbieski, Helmut Fuchs, Martin Hrabé de Angelis, Mitul Desai, Sarah Bricault, Alan Jasanoff, Vasilis Ntziachristos, Jüergen Plitzko, Gil G. Westmeyer

## Abstract

We have genetically controlled compartmentalization in eukaryotic cells by heterologous expression of bacterial encapsulin shell and cargo proteins to engineer enclosed enzymatic reactions and size-controlled metal biomineralization. The orthogonal shell protein (EncA) from *M. xanthus* efficiently auto-assembled inside mammalian cells into nanocompartments to which sets of native (EncB,C,D) and engineered cargo proteins self-targeted. This enabled localized bimolecular fluorescence and enzyme complementation with selective access to substrates via the pores in the nanoshell. Encapsulation of the enzyme tyrosinase lead to the confinement of toxic melanin production for robust detection via multispectral optoacoustic tomography (MSOT). Co-expression of ferritin-like native cargo (EncB or EncC) resulted in efficient iron sequestration that produced substantial contrast by magnetic resonance imaging (MRI) and enabled magnetic cell sorting. The monodisperse, spherical, and iron-loading nanoshells also proved to be excellent genetically encoded markers for cryo-electron tomography (cryo-ET). In general, eukaryotically expressed encapsulins enable cellular engineering of spatially confined multicomponent processes with versatile applications in multiscale molecular imaging, as well as intriguing implications for metabolic engineering and cellular therapy.

---

Compartmentalization, the spatial separation of processes into closed subspaces, is an important principle that has evolved on several biological scales: multi-enzyme complexes that channel substrates, nanocompartments built entirely from proteins, as well as membrane-enclosed organelles, cells, and organs. Compartments make it possible to generate and maintain specific local conditions that can facilitate interactions and reactions in confined environments^1^, such that they can isolate toxic reaction products, protect labile intermediate products from degradation, or separate anabolic from catabolic processes^2^. Whereas eukaryotes possess many membrane-enclosed organelles, membranous compartments are not known in bacteria with a notable exception of magnetosomes in magnetotactic bacteria, in which specific reaction conditions are maintained that enable magnetic biomineralization^3–4^. However, nanocompartment shells built entirely from protein complexes can serve functions in prokaryotes that are analogous to eukaryotic organelles^5^.

Intense work has been invested in engineering compartments in prokaryotic systems and yeast to realize features such as substrate channeling for biotechnological production processes^1,6,7^. In contrast, no orthogonal compartments with self-targeting cargo molecules exist to date for use in mammalian cells. Such a system could for instance enable cellular engineering of reaction chambers that would endow genetically modified mammalian cells with new metabolic pathways that may include labile intermediate products or spatially confined toxic compounds. Engineered orthogonal compartments in eukaryotic cells may also enable size-constrained synthesis of biomaterials via, *e.g*., metal biomineralization processes occurring under specific localized environmental conditions.

With regards to protein complexes as building blocks for addressable nanocompartments, viruses and virus-like particles have been expressed in bacterial hosts to encase fluorescent proteins^8–12^, enzymes^13–16^ and even multi-enzymatic processes^17,18^. In mammalian systems, vault proteins (vaults) have been explored, which are ribonucleoprotein complexes enclosed by ~60 nm large envelope structures^19^ into which foreign cargo proteins such as fluorescent proteins or enzymes can be packaged^20–22^. However, vaults have openings on both ends and are endogenously expressed by many eukaryotic cells such that overexpressing them may interfere with their natural functions^23^. With respect to protein shell structures that can incorporate iron, the central iron storage protein ferritin has been overexpressed to generate MRI contrast under certain conditions although the core size is only ~6 nm containing only ~2000 iron atoms on average per core, which can result in only low magnetization^24–26^. Viral capsids such as the ones from cowpea chlorotic mottle virus (CCMV) have also been equipped with iron-binding sites that lead to accumulation of iron. Expression, assembly, and iron loading in mammalian cells, however, has not yet been demonstrated^27^.

In search of a versatile nanocompartment-cargo system for heterologous expression in eukaryotic cells, we were intrigued by the recently discovered class of prokaryotic proteinaceous shell proteins called encapsulins because they possess a set of very attractive features: (1) A single shell protein - without the need for proteolytic processing - is sufficient to form comparably large shell-like architectures (~18 nm or ~32 nm) auto-assembled from 60, or 180 identical subunits with a triangulation number (T) of one (T=1) or three (T=3), respectively^28^. (2) The assembled shells are pH-resistant and temperature-stable^29^. (3) A versatile set of native cargo molecules including enzymes exist that are packaged into shell structures via specific encapsulation signals defined by a short terminal peptide sequence^29^. (4) The pore size of ~5 Å allows the shell architecture to channel small molecular substrates^30^. (5) *M. xanthus* encapsulin was also shown to use ferritin-like cargo proteins to import and sequester iron inside the nanoshell. These proteins have an “open ferritin structure” and possess ferroxidase activity^28,30^. A model based on structural data from *T. maritima* encapsulins (with T=1) assumes that the ferritin-like protein docks into the shell where it obtains ferrous iron through the pores which it then oxidizes for deposition of up to an estimated 30,000 iron atoms per shell in the case of *M. xanthus*^28,30,31^. This is an order of magnitude more than can be contained inside a ferritin core expressed in eukaryotic cells. (6) The termini of the shell protein extend to the inner and outer surface, respectively, such that surface functionalizations are possible. The outer surface can for instance be functionalized to install specific targeting moieties^32–34^. Encapsulin variants were also purified when secreted from HEK293 cells to present glycosylated epitopes for an innovative vaccination approach^35^. Recently, the inner surface of the shell from *T. maritima* was also modified with silver-binding peptides to cause local silver precipitation in *E. coli* but the ferritin-like cargo was deleted to achieve this feature^36^. (7) Non-native cargo proteins including enzymes can be addressed to the inside of the nanocompartment via a short encapsulation signal^11–37^.

This excellent set of studies showed the feasibility and utility of biotechnological production of encapsulins as biomolecular scaffolds and targetable vehicles and probes.

## RESULTS

We here introduce engineered encapsulins modified from *M. xanthus* in the context of genetic programming of orthogonal and addressable cellular compartments in mammalian cells.

We specifically demonstrate that eukaryotically expressed encapsulins not only auto-assemble at high density and without toxic effects but that self-targeting and encapsulation of cargo molecules still efficiently occur in mammalian cells.

We furthermore show localized enzymatic reactions in the nanocompartment useful for optical and optoacoustic imaging, as well as confined iron accumulation within the nanocompartments that labels cells for detection by MRI and cryo-electron tomography (cryo-ET), demonstrating the potential of encapsulins as genetic markers across modalities. In addition, the iron-sequestration inside the nanoshells affords magnetic manipulation of cells genetically labelled with encapsulins.

### Encapsulin expression and auto-assembly

Based on the favorable set of features introduced above, we chose to heterologously overexpress the encapsulin shell protein from *M. xanthus* in HEK293T cells. We tagged the nanoshell with an outward facing FLAG epitope (A^FLAG^) and found it to express strongly without and with the native cargo molecules from *M. xanthus*, denoted encapsulins B,C, and D^28^.

Co-expression of Myc-tagged B, C, or D alone, or a combination of all three non-tagged proteins (via co-transfection or a P2A construct, **Fig. 1b**), co-immunoprecipitated with A^FLAG^ as visualized on silver-stained SDS-PAGE (**Fig. 1c** middle panel). A corresponding Western Blot against the FLAG (**Fig. 1c** upper panel) or Myc-epitope (**Fig. 1c** lower panel) confirmed the identities of the protein bands (A^FLAG^: 32.9 kDa, ^Myc^B: 18.5 kDa, ^Myc^C: 15,4 kDa, ^Myc^D: 12.5 kDa).

**Figure 1:**
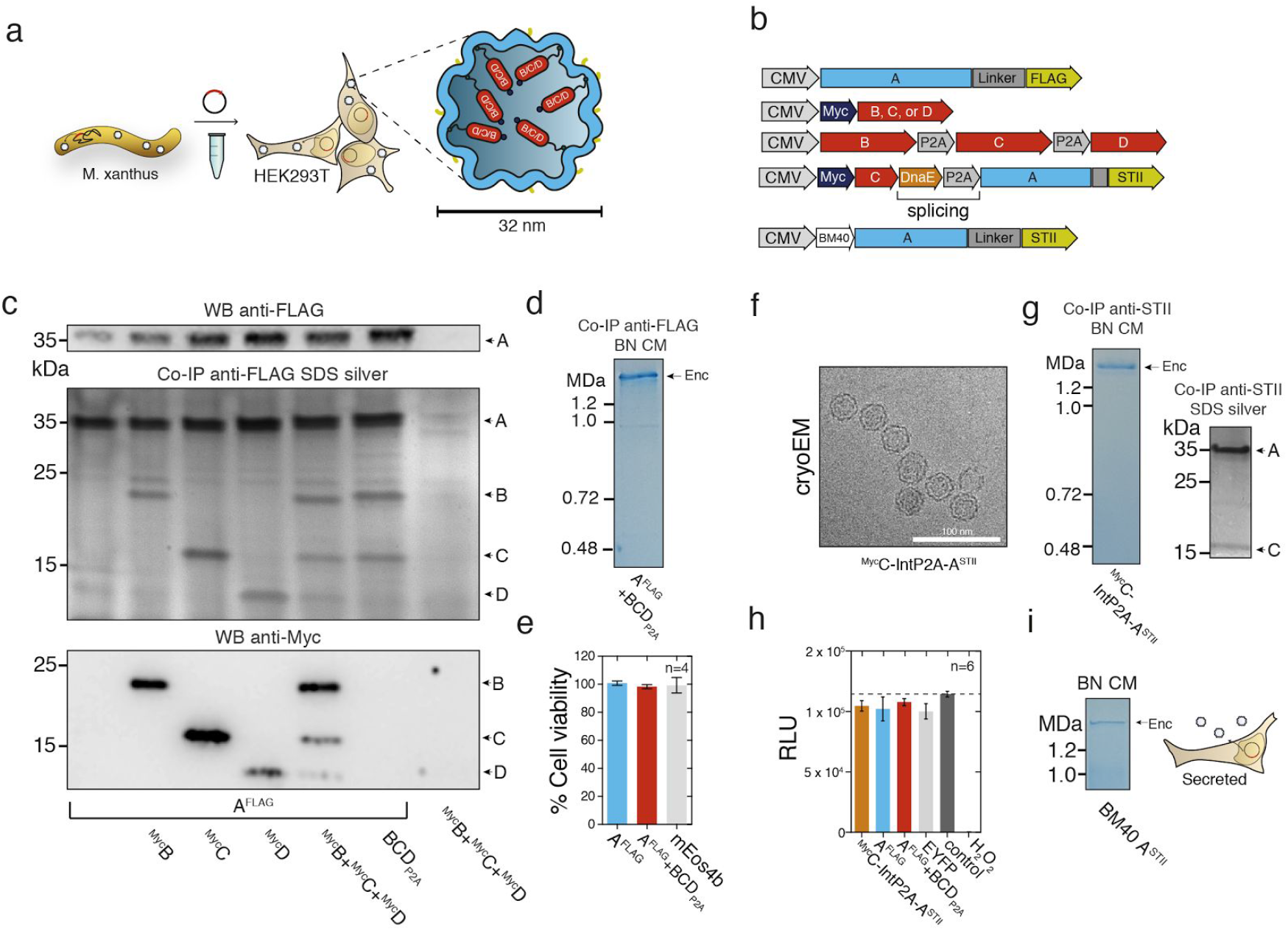
Assembly of eukaryotically expressed encapsulin nanocompartments and targeting of native cargo proteins in HEK293T cells. **(a)** Schematic of the heterologous expression of surface-modified encapsulin variants loaded with endogenous cargo proteins. **(b)** Genetic constructs encoding the shell protein A (light blue) with a FLAG-tag as C-terminal surface modification as well as individual Myc-tagged cargo proteins (red) B, C, and D that can also be combined in a multi-gene expression construct (BCD_P2A_). Also shown is a P2A bicistronic expression construct encoding StrepII-tagged (^STII^) nanocompartments containing Myc-tagged C as cargo protein (^Myc^C-IntP2A-A^STII^) as well as a variant with a N-terminal BM40 secretion peptide and StrepTagII (^STII^). **(c)** Co-immunoprecipitation of A^FLAG^ and silver-stained SDS-PAGE from cells co-expressing just B, C, or D, or a combination of these three proteins expressed either via a mixture of individual DNA constructs (^M^y^c^B+^Myc^C+^Myc^D), or by a multi-gene expression construct (BCD_P2A_). The top panel shows a Western Blot (WB) against the exterior FLAG-Tag in A^FLAG^. The bottom panel shows the corresponding WB against the Myc epitope. **(d)** Coomassie-stained Blue Native PAGE (BN CM) of purified material from HEK293T expressing A^FLAG^ and BCD_P2A_ yielding a band above 1.2 MDa. **(e)** Cell viability after 48 h of overexpression of encapsulins (A^FLAG^) with or without cargo (BCD_P2A_) assessed by a LDH release assay. A construct expressing the fluorescent protein mEos4b served as a control. (The bars represent the mean ± SEM, *n*=4, Kruskal-Wallis, *p*=0.1965, No significant differences in Dunn’s nonparametric comparison between mEos4b and A^FLAG^ with or w/o BCD_P2A_ at α=0.05). **(f)** Cryo-electron microscopy image of material from HEK293T cells expressing ^Myc^C-IntP2A-A^STII^ purified via Strep-tag II/Strep-Tactin XT affinity chromatography showed the assembled nanospheres of ~32 nm diameter. Scale bar is 100 nm. **(g)** The corresponding BN-PAGE analysis of the identical material revealed a single band larger than 1.2 MDa and the accompanying silver-stained SDS-PAGE showed the coprecipitation of the cargo ^Myc^C with the StrepTagII-modified nanoshell. **(h)** Luciferase-based cell viability assay after 48 hours of overexpression of ^Myc^C-IntP2A-A^STII^ and A^FLAG^ with or without cargo BCD_P2A_. Cells overexpressing the fluorescent protein EYFP as well as untransfected HEK293T cells served as a negative controls. To induce toxicity as positive control, un-transfected HEK293T cells were treated with 1 mM H_2_O_2_ 24 h prior to the assay (The bars represent the mean ± SEM, *n*=6, Kruskal-Wallis, *p*=0.0041, no significant differences in Dunn’s nonparametric comparison between A^FLAG^ with and w/o BCD_P2A_ or ^Myc^C-IntP2A-A^STII^ vs. EFYP or control at α=0.05). **(i)** BN CM loaded with cell culture supernatant of HEK293T cells expressing A^STII^ with an N-terminal BM40 secretion signal showed a single band >1.2 MDa.

Furthermore, a corresponding Blue Native PAGE (BN-PAGE) of immunoprecipitated FLAG-tagged material from cells expressing A^FLAG^ together with BCD_P2A_ revealed a band with apparent molecular weight of above 1.2 MDa indicating autoassembly of encapsulin protein complexes and self-targeting of all native cargo proteins (**Fig. 1d**).

The strong expression of A^FLAG^ without or with loaded cargo did not result in a reduction of cell viability when compared to cells overexpressing a fluorescent protein as assessed by a viability assay based on lactate dehydrogenase (LDH) release (**Fig. 1e**). We also generated a StrepTagII-labelled variant of the shell that allowed for convenient Strep-Tactin affinity chromatography. We co-expressed this version of the nanosphere with Myc-tagged C as cargo protein (^Myc^C-IntP2A-A^STII^, **Fig. 1b**). Purified material from HEK293T cells analyzed by cryo-electron tomography showed assembled nanospheres of 32.4 ± 1.7 nm (**Fig. 1f**) corresponding to the single band >1.2 MDa in size on BN-PAGE (**Fig. 1g**). Again, no effect on cell viability was detected for this construct tested by a luciferase-based viability assay and compared to A^FLAG^ with and without cargo (BCD_P2A_), as well as negative (1 mM H_2_O_2_ added 24 h prior to the assay) and positive controls (EYFP and untransfected HEK293T) (**Fig. 1h**). Furthermore, robust secretion of StrepTagII-modified encapsulins from HEK293T cells was possible by addition of a N-terminal BM40 secretion signal as shown by Coomassie-stained BN-PAGE of material present in the cell culture supernatant (**Fig. 1i**).

### *In vivo* expression of encapsulins

To achieve *in vivo* expression of encapsulins, we generated a coexpression construct that encoded both the nanoshell A^FLAG^ and the ferritin-like protein B from a single plasmid that was small enough to be packaged into an Adeno-associated virus (AAV) (**Supplementary Fig. 1a**). After transduction of murine brains via intracranial injections of this viral vector co-expressing A^FLAG^ and B^M7^ via a P2A peptide, we observed robust neuronal expression of the shell protein (**Supplementary Fig. 1d**). Silver-stained BN-PAGE and SDS-PAGE of immunoprecipitated (anti-FLAG) proteins extracted from murine brain showed that the nanocompartments assembled *in vivo* and that the cargo B^M7^ was associated with the shell (**Supplementary Fig. 1e**). Similar *in vivo* results could be obtained by expressing the nanoshell and ferritin-like B cargo via an IRES site (**Supplementary Fig. 1f**).

### Simultaneous encapsulation of sets of engineered cargo molecules for bimolecular fluorescence complementation

Next, we sought to test whether engineered cargo proteins could be efficiently targeted to the nanocompartment. To accomplish this, we either C-terminally appended a minimal encapsulation tag^29^ that we found to only necessitate 8 amino acids (EncTag) to different reporters, or fused the reporter protein to the C-terminus of either B or C (**Fig. 2a**). Via the EncTag, the photoactivatable fluorescent protein mEos4b^38^ was targeted to the nanocapsules as shown by co-immunoprecipitation (Co-IP) and fluorescence imaging of the encapsulin band on a BN gel (**Fig. 2b**). To demonstrate that more than one engineered cargo molecule could be targeted to the nanocompartment simultaneously, we fused the two halves of split PAmCherry1 (PA-s1, PA-s2) to either B or C (B-PA-s2: 27.0 kDa, C-PA-s1: 33.1 kDa) and tested for bimolecular fluorescence complementation (BiFC) within the nanocompartment^39^. Either of these components could be co-immunoprecipitated with A^FLAG^ as shown by silver-stained SDS-PAGE (**Fig. 2c**). The photoactivation of the complemented split PAmCherry1 inside the encapsulins could also be detected via fluorescence imaging of the corresponding BN-PAGE (**Fig. 2c** right panel). Co-expression of both split halves together with A^FLAG^ lead to a strong increase of photoactivatable fluorescent signal throughout the cytosol of HEK293T cells as quantified by confocal microscopy compared to cells that did not express A^FLAG^ (**Fig. 2d**). In addition to controlling and restricting the localization of cargo molecules to the encapsulin lumen by targeting two split parts that only effectively reconstitute a functional unit inside the encapsulin, we present an alternative by fusing cargo proteins to a FKBP12-derived destabilizing domain (DD) that yields rapid proteasomal degradation of the cargo if not protected inside the encapsulin or stabilized by a synthetic small molecule^40^. We show that co-expressing A^FLAG^ + DD-mEos4b-EncTag in HEK293T yielded significantly higher mean fluorescence values than DD-mEos4b-EncTag alone, indicating that cargos inside the encapsulin are protected from proteolytic degradation (**Supplementary Fig. 2b, c**).

**Figure 2:**
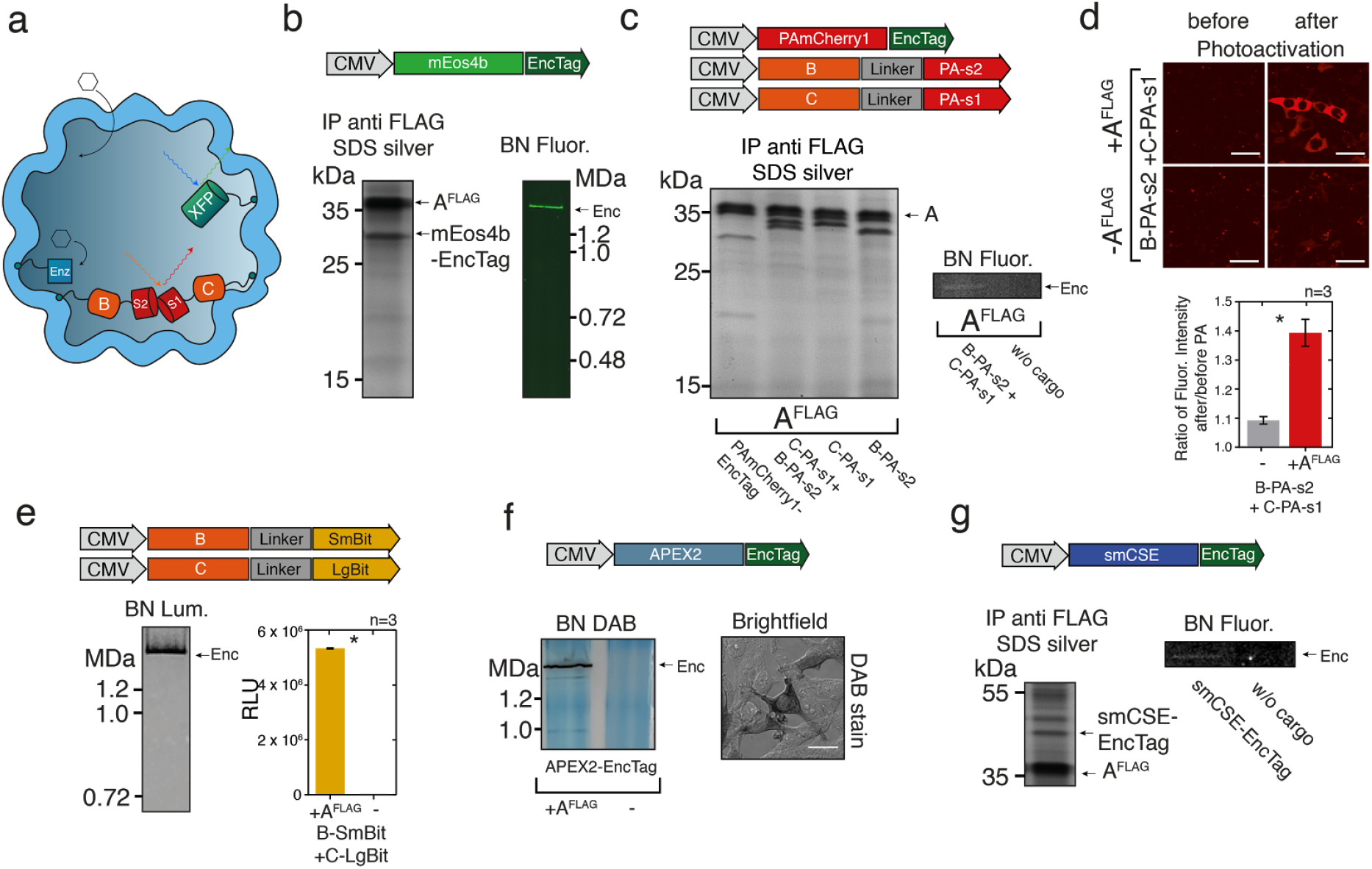
Modified encapsulin nanocompartments can host sets of engineered cargo proteins. **(a)** Overview schematic of sets of cargo molecules for bimolecular fluorescence and enzyme complementation inside the nanocompartment. Targeting of foreign cargo proteins can be achieved either via a minimal C-terminal encapsulation signal (EncTag) or via C-terminal fusions to the native cargo proteins B,C or D. **(b)** Cargo loading of the photoactivatable fluorescent protein mEos4b C-terminally modified with an EncTag into the nanocompartment composed of A^FLAG^ was demonstrated by co-immunoprecipitation (Co-IP) against the FLAG epitope followed by silver-stained SDS-PAGE (left panel). The right panel shows the fluorescent Enc band on a BN-PAGE loaded with whole cell lysate visualized on an UV-imager. **(c)** Silver-stained SDS-PAGE from a co-immunoprecipitation (Co-IP) of A^FLAG^ co-expressed with photoactivatable mCherry1 with EncTag (PAmCherry1-EncTag) or with either one of the halves of split PAmCherry1 fused to C or B, or a combination of both (C-PA-s1+B-PA-s2). Fluorescence originating from complemented split PAmCherry1 inside the encapsulins was detected on BN-PAGE loaded with whole cell lysates of cells expressing A^FLAG^ and C-PA-s1+B-PA-s2 after 2 min of PA on an UV-imager. **(d)** Live cell confocal microscopy images (scale bar: 20 μm) of HEK293T cells expressing B-PAs1 and C-PAs2 with or without the shell-protein A^FLAG^ before and after 60 seconds of photoactivation (PA) with 405 nm (upper panel) demonstrating efficient bimolecular fluorescence complementation inside encapsulin compartments. Fluorescence of photoactivated split PAmCherry1 was excited using a 561 nm laser. Fluorescence signals of the sample without and with A^FLAG^ were quantified by calculating the ratio of the mean signal after PA divided by the signal before PA. (The bars represent the mean ± SEM, *n*=3, *p*=0.0123, unpaired t-test) (lower panel). **(e)** Luminescence signal from BN-PAGE incubated with luciferase substrate and loaded with whole-cell lysates of HEK293T co-expressing split luciferase fragments fused to either B or C (B-SmBit, C-LgBit) and A^FLAG^ (left panel). The luminescent band corresponds to the complemented split luciferase inside the assembled nanocompartment. The bar graph shows the corresponding total luminescence signals from the cell lysates expressing B-SmBit and C-LgBit with or without A^FLAG^, (mean ± SEM, *n*=3, *p*<0.0001, unpaired t-test) (right panel). **(f)** DAB-stained BN-PAGE of whole cell lysates of HEK293T cells expressing APEX2-EncTag with or without A^FLAG^ demonstrating peroxidase activity of APEX2 inside the encapsulin compartment. The brightfield microscopy image (lower panel) shows stained HEK293T cells co-expressing A^FLAG^ and APEX2-EncTag incubated with 0.7 mg/ml DAB and H_2_O_2_ in 60 mM Tris buffer for 2 min. Scale bar:20 μm. **(g)** Co-IP of the putative cystathionine γ-lyase (SmCSE-EncTag) with encapsulin analyzed by silver-stained SDS-PAGE. The right panel shows a BN gel of encapsulated SmCSE-EncTag with on-gel formation of CdS nanodots after supplementation of 0.5 mM cadmium acetate and 4 mM L-cysteine for 2 h visualized by UV-fluorescence.

### Compartmentalized enzymatic reactions

To showcase the ability to utilize the eukaryotically expressed encapsulins as bioengineered reaction chambers with pores that can constrain passage of reactants and reaction products, we targeted several enzymes to the nanocapsules. In the presence of A^FLAG^, the split luciferase^41^ parts LgBit and SmBit fused to C and B (C-LgBit: 32.7 kDa, B-SmBit: 19.6 kDa) were complemented to functional enzymes as demonstrated by bioluminescence detection from BN-PAGE (**Fig. 2e** left gel) and from total lysate (**Fig. 2e** bar graph).

Similarly, we showed that the engineered peroxidase APEX2^42^ can polymerize Diaminobenzidine (DAB) when targeted to the nanocompartment (APEX2-EncTag; 31.0 kDa) as indicated by the generation of photoabsorbing DAB polymers associated with the BN-PAGE band corresponding to the assembled nanosphere (**Fig. 2f**). Since the electrophoretic mobility of protein complexes on BN-PAGE also depends on their hydrodynamic size and shape^43^, cargo-loading can be confirmed by observing an identical migration behavior of loaded as compared to unloaded capsules (**Supplementary Fig. 3**).

Moreover, we targeted the putative cystathionine γ-lyase (SmCSE) to the nanospheres via an EncTag (SmCSE-EncTag: 43.9 kDa) as shown by Co-IP with A^FLAG^. In the presence of L-cysteine, this enzyme was reported to catalyze a conversion of cadmium acetate in aqueous solution into Cadmium sulfide (CdS) nanocrystals such that they would generate a photoluminescence signal under UV illumination characteristic for crystal formation at quantum confined sizes^44^. Interestingly, we could detect a photoluminescence signal from the BN-PAGE band corresponding to encapsulin loaded with SmCSE-EncTag after on-gel incubation with cadmium acetate and L-cysteine indicating that the SmCSE-EncTag cargo was enzymatically active when bound into the shell (**Fig. 2g**).

### Bioengineered melanosomes as non-toxic gene reporters for optoacoustics

We next sought to utilize selective passage of small substrates through the nanoshell to load the compartments with tyrosinase as cargo which is the sole enzyme generating the strongly photoabsorbing polymer melanin from the amino acid tyrosine. Because of these attractive features, tyrosinase has been used as a gene reporter for optoacoustic tomography^45,46^, an imaging modality that maps the distribution of photoabsorbing molecules in tissue by locating the ultrasonic waves that they emit in response to local heating upon laser absorption^47,48^. However, melanin production is toxic to cells if not confined in melanosomes, membraneous compartments of specialized cells^49,50^. We thus chose a soluble tyrosinase from *Bacillus megaterium*^51^ that we thought could still be functional as a fusion protein to the native cargo D (^Myc^D-BmTyr: 47.7 kDa) serving as targeting moiety. Indeed we could observe generation of melanin on the BN-PAGE band corresponding to the assembled nanocompartment (**Fig. 3b**). In cells expressing the encapsulin-targeted tyrosinase and the shell A^STII^, we observed strong melanin formation by brightfield microscopy without the strong toxicity apparent morphologically in control cells expressing just the tyrosinase (**Fig. 3c**, white arrows). Encapsulation of the tyrosinase also lead to a significant reduction of cell viability as assessed by a luciferase-based viability assay (**Fig. 3d**). Cells expressing melanin-filled encapsulins were dark in color (**Fig. 3e inset**) and thus generated strong photoacoustic signal even when referenced against strongly absorbing synthetic ink (optical density of 0.2) (**Fig. 3e**).

**Figure 3:**
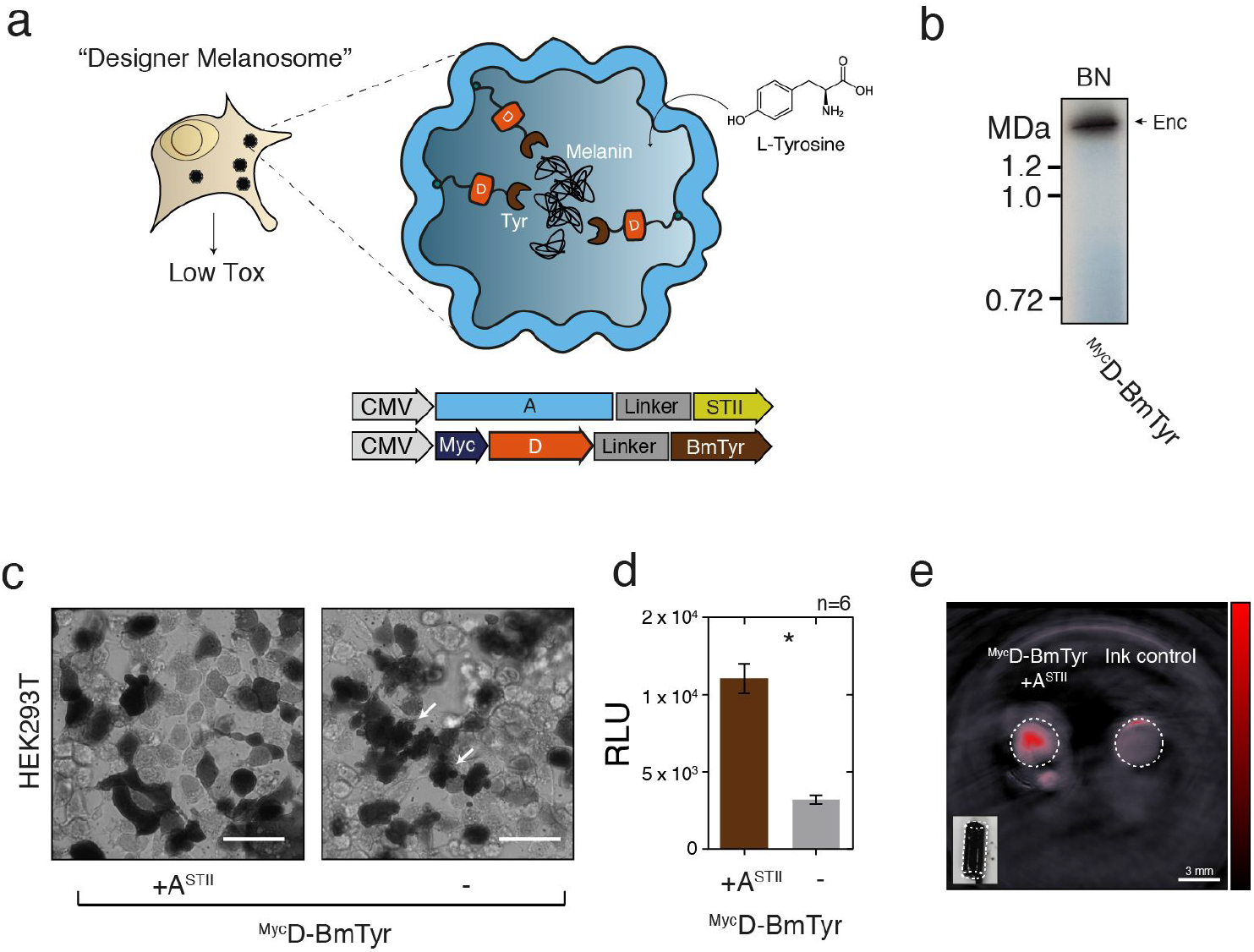
Bioengineering of a melanosome by targeting melanin-generating tyrosinase to the encapsulin compartment. **(a)** Schematic of the detoxifying effects of compartmentalized melanin-production by encapsulated tyrosinase from *Bacillus megaterium* targeted to the nanocompartment via a fusion to the native cargo D. The substrate L-tyrosine enters the compartment via the pores in the nanoshell. **(b)** BN-PAGE showing on-gel production of melanin tyrosinase expressed in HEK293T cells fused to Myc-tagged encapsulin-cargo D(^Myc^D-BmTyr) to encapsulate it in the assembled nanoshell. Brown colorization of the band was observed after incubation with 2 mM L-tyrosine and 100 μM CuCl_2_ in PBS (pH 7.4) for 1 hour at 37°C. **(c)** Brightfield images of HEK293T cells expressing ^Myc^D-BmTyr with and without StrepTagII-modified shell (A^STII^) after 48 h of expression. 24 h post transfection cells were supplemented with 1 mM L-tyrosine and 10 μM CuCl_2_. Cell protrusions (white arrows) were apparent indicating toxic effects of overexpression of non-encapsulated tyrosinase. Scale bar: 20 μm. **(d)** Corresponding luciferase based viability assay of HEK293T cells treated as in subpanel c overexpressing ^Myc^D-BmTyr with or without A^STII^ after 48 h. (The bars represent the mean ± SEM, *n*=6, *p*<0.0001, unpaired t-test) **(e)** Images of of two tubular phantoms (transversal slice) obtained by multispectral optoacoustic tomography (MSOT). The phantoms were filled with ~10^7^ cells in 1.5 % low melting agar expressing ^Myc^D-BmTyr with A^STII^ (supplementation as in **c** and **d**) or containing highly concentrated ink (OD=0.2) as control showing the strong contrast obtained between 690 and 900 nm from the melanin-filled encapsulins. The coefficients obtained from linear unmixing of the optoacoustic spectra with a melanin reference spectrum are displayed on the red colormap overlaid on the image obtained at 720 nm. The lower left inset shows a color photograph of the tubular phantom containing the cells. Scale bar: 3 mm.

### Size-controlled iron biomineralization

Another important reason that guided our selection of encapsulins from *M. xanthus*, was a report showing that they can deposit iron via the ferritin-like cargo B and C into relatively large compartments (~32 nm, T=3)^28^. We thus investigated whether this functionality could also be realized in eukaryotic cells to enable spatially confined iron deposition sequestered away from the complex signaling network controlling mammalian iron homeostasis. We thus generated a stable cell line co-expressing the nanoshell (A^FLAG^) with all native cargo proteins (B, C, D) via a dual-promoter construct (Fig. 4a) and observed long-term and robust expression of all components shown by co-immunoprecipitation with A^FLAG^ (Fig. 4b, left panel). Transient co-expression of the ferrous iron transporter MmZip14^FLAG^ (Zip14) in the stable cell line resulted in a strong dose-dependent iron loading at ferrous ammonium sulfate (FAS) concentrations between 0.25–1.25 mM already after 48 h of supplementation as detected on BN-PAGE via DAB-enhanced Prussian Blue staining (DAB PB) (**Fig. 4b** right panel).

**Figure 4:**
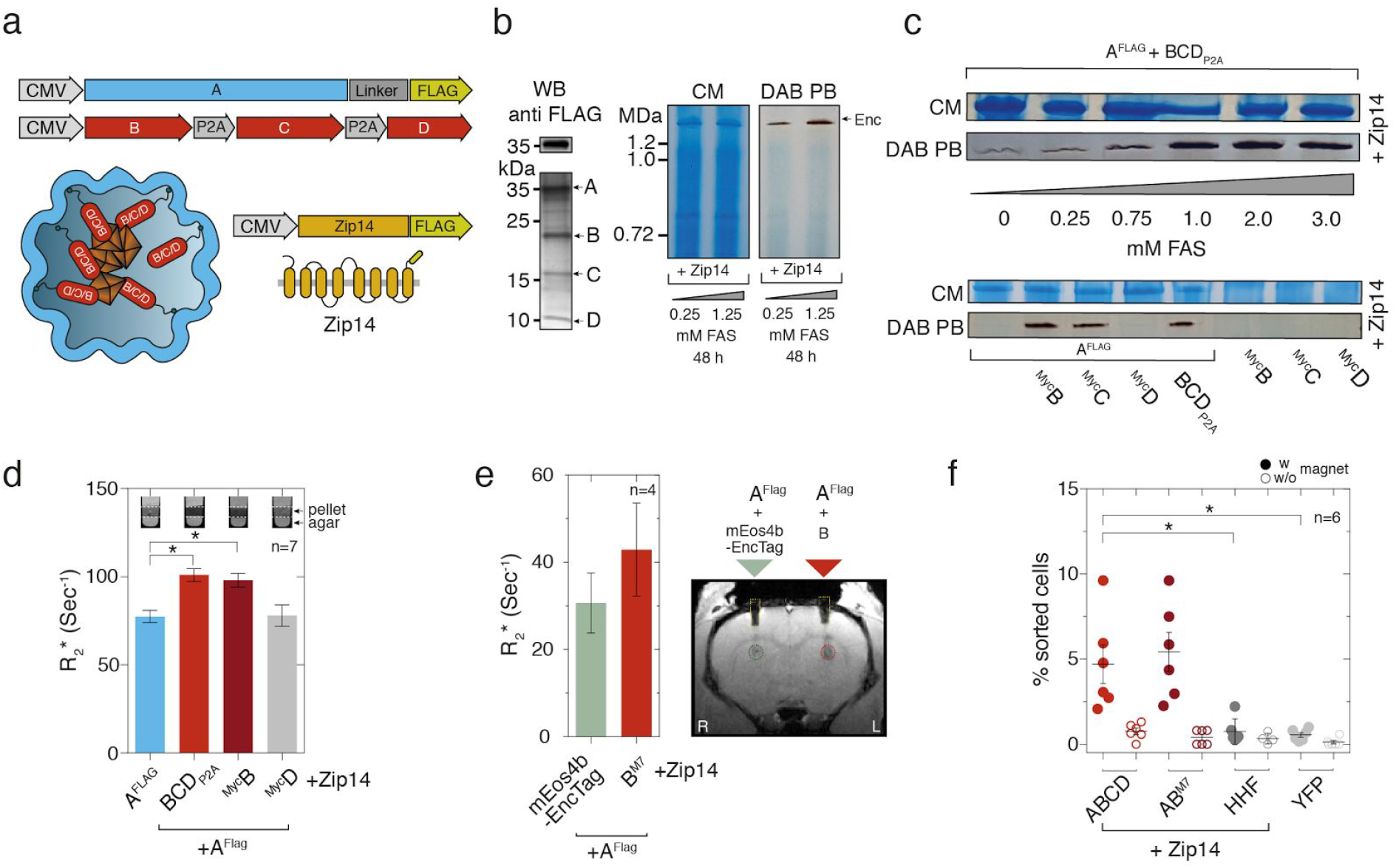
Efficient iron-loading of eukaryotically expressed encapsulin nanospheres enabled detection by MRI and magnetic sorting. **(a)** Schematic of a dual-promoter construct used for generation of a stable cell line expressing A^FLAG^ and all native cargos B, C, and D. Also depicted is a construct encoding the iron-transporter MmZip14^FLAG^ (Zip14) used to transport additional amounts of iron into the cell. **(b)** Co-immunoprecipitation (Co-IP) against the FLAG epitope from a whole cell lysate of a stable HEK293T clone expressing A^FLAG^ together with B, C, and D analyzed by silver-stained SDS PAGE and the corresponding WB against the FLAG epitope (left panel). The pair of Blue Native (BN) gels visualize proteins from whole cell lysates via Coomassie staining (CM) (left panel) and iron content via treatment with DAB enhanced Prussian Blue (DAB PB) (right panel) from the same stable cell line. Robust iron-loading of the assembled nanocompartments was achieved by transient co-expression of MmZip14^FLAG^-IRES-ZsGreen1 in which case 0.25 mM ferrous ammonium sulfate (FAS) for 48 hours was sufficient to see strong iron loading. **(c)** BN gel stained with CM or DAB PB loaded with whole cell lysates of HEK293T cells transiently expressing A^FLAG^ + BCD_P2A_ and Zip14^FLAG^ supplemented with different concentrations of FAS (0–3 mM) for 48 h (upper panel). The strong bands, which correspond to the assembled nanoshell, indicate high expression levels of encapsulins and efficient, dose-dependent iron loading. The lower panel shows a CM and DAB PB-stained BN gel from whole cell lysates of HEK293T cells expressing Zip14^FLAG^ and different combinations of native cargo molecules: ^Myc^B, ^Myc^C and ^Myc^D alone, or all three (BCD_P2A_) with or without A^FLAG^. The robust DAB PB stains show that the ferritin-like cargo proteins B or C are sufficient for efficient iron loading into encapsulins. FAS was supplemented at 2.5 mM for 48 h. **(d)** Relaxometry measurements by magnetic resonance imaging (MRI) conducted on cell pellets (~10^7^ cells) from HEK293T cells transiently expressing A^FLAG^ + BCD_P2A_, ^Myc^B, or ^Myc^D, or A^FLAG^ alone (1 mM FAS supplementation for 24 h with expression of Zip14^FLAG^). Expression of A^FLAG^ + BCDp_2_A or ^Myc^B shows a significantly enhanced R_2_*-relaxation rate as compared with the nanocompartment alone or loaded with ^Myc^D; ferritin-like cargo B was sufficient to generate and increase in R_2_* in the presence of the A^FLAG^ nanocompartment. (The bars represent the mean ± SEM, *n*=7 from 4 independent experiments, Kruskal-Wallis, *p*=0.0047, stars indicate significance from Dunn’s nonparametric comparison vs. A^FLAG^ at α=0.05). The insets show MRI slices (13.5 ms echo time) through test tubes in which cells were pelleted on a layer of agar. **(e)** *In vivo* MRI detection of HEK293T cells transiently co-expressing A^FLAG^ together with ferritin-like B^M7^ that were xenografted into rat brains. As compared to cells co-expressing A^FLAG^ together with the fluorescent protein mEos4b-EncTag as control cargo, there was a trend (The bars show the mean ± SEM, *p* =0.0834, paired t-test, *n*=4) towards elevated transverse relaxation rates measured at the injection site of A^FLAG^+B^M7^ expressing cells 24 h post injection. The image on the right shows a coronal MRI slice through a rat brain (gradient echo, 5 ms echo time) indicating the regions of interest (ROIs) defined over the injection sites. The low MRI signals in the two cortical regions (yellow dashed lines) and above are due to the implanted plastic guide cannulae and head post (please see Online Methods). **(f)** Magnetic sorting of HEK293T cells co-expressing A^FLAG^ and BCD_P2A_, A^FLAG^ and B^M7^, or human H-chain ferritin (HHF) with Zip14^FLAG^ and treated with 2.5 mM FAS for 48 h. Control cells were only expressing EYFP. Independent samples were sorted on columns inside and outside a magnetic field to control for unspecific retention in the mesh of the column (The horizontal lines represent the mean ± SEM, *n*=6 from 3 independent experiments, Kruskal-Wallis, *p* =0.0007, stars indicate significance from Dunn’s nonparametric comparisons at α=0.05).

Efficient iron-loading could also be achieved by transient expression of A^FLAG^ + BCD_P2A_ together with Zip14. Under these conditions, iron supplementation with ~0–3 mM FAS for 48 hours led to a strong dose-dependent iron loading of the nanocompartment that saturated at ~1 mM FAS as shown by Coomassie and DAB-enhanced Prussian Blue BN PAGE (**Fig. 4c**, upper panel). Interestingly, when we tested the cargo molecules individually for their ability to load iron into the nanosphere, we found that co-expression of only B or C generated equally strong DAB PB bands as compared to BCD_P2A_, indicating that B or C are sufficient for iron-deposition inside the nanocompartment.

In contrast, co-expression of D with A^FLAG^ or any of the cargo molecules without presence of A^FLAG^ did not lead to discernable DAB PB signals (**Fig. 4c**, lower panel). In a cell viability assay, we found no impairment of the cells when Zip14 was co-expressed together with A^FLAG^ and BCD_P2A_ or just B. However, ~7% of cells showed reduced viability when BCD_P2A_ were expressed without the nanocompartment (Mann Whitney test, p=0.0238) or when just the fluorescent protein mEos4b-EncTag was expressed (**Supplementary Fig. 4b**) indicating that in the absence of the nancompartment the imported iron was not sufficiently sequestered by the endogenous iron homeostasis machinery.

We furthermore tested variants of A with N-terminal fusions with peptide sequences from *Magnetospirillum magneticum* Mms (6 and 7) proteins reported to aid in templating iron mineralization^52^ but found no additional benefit of these modified inner surfaces over A^FLAG^. In addition, we analyzed several variants of the cargo proteins B and C, fused C-terminally to peptides from Mms proteins (superscripts M6, M7, please see supplemental methods). These data confirmed that either B or C are sufficient to load the nanocompartment with iron and showed that no obvious additional iron loading resulted from the presence of the Mms peptides (**Supplementary Fig. 4d**).

### Encapsulins enable detection via MRI and Magnetic Sorting

Next, we were interested in whether the strong iron accumulation inside eukaryotically expressed encapsulin shells would yield significant contrast by MRI. We thus expressed A^FLAG^ alone or together with either all native cargos BCD_P2A_ or just ^Myc^B, or ^Myc^D and Zip14 and subjected cell pellets to relaxometry measurements by MRI. The nanocompartment A^FLAG^ co-expressed with all native cargo or just with ferritin-like B lead to a similar increase in R_2_*-relaxation rates as compared to just A^FLAG^ (**Fig. 4d**, *n* =7 from 4 independent experiments, Kruskal-Wallis, *p*=0.0047, Dunn’s nonparametric comparison vs. A^FLAG^ at α=0.05). This indicated again that co-expression of B was sufficient to generate efficient iron deposition inside the nanoshell.

We subsequently sought to test whether cells genetically labelled with encapsulins could be detected by MRI *in vivo*. As an initial assessment, we thus xenografted cells co-expressing A^FLAG^ together with B^M7^ into rat brains and obtained R_2_*-relaxation maps that showed a trend towards elevated relaxation rates (*p*=0.0834, paired t-test, *n*=4) at the injection site as compared to xenografted cells in which the fluorescent protein mEos4b-EncTag was used as a control cargo (**Fig. 4e**).

In addition to MRI contrast, the strong iron accumulation also allowed us to magnetically sort cells expressing A^FLAG^ + BCD_P2A_ or B^M7^ at significantly higher percentages than when human H-chain ferritin was expressed or just yellow fluorescent protein (EYFP) (Kruskal-Wallis, *p* = 0.0007, Dunn’s nonparametric comparisons at α=0.05, *n* =6 from 3 independent experiments, **Fig. 4f**).

### Encapsulins as genetically encoded markers for electron microscopy

Given that the iron loading of the encapsulins was very efficient and observable at the population level, we next assessed how well individual nanocompartments could be detected via electron microscopy in cells such that they could be used as genetically encoded markers. We thus grew HEK293T cells stably expressing the shell protein A^FLAG^ and BCD_P2A_ using a dual promoter vector on a transmission electron microscopy (TEM) grid, vitrified them by plunge-freezing and produced lamellae by cryo-focused ion beam (cryo-FIB) milling for cellular cryo-electron tomography (cryo-ET). The heterologously expressed encapsulins were readily detected as clearly discernable nanospheres (**Figure 5a, Supplementary Figure 5 a,b**) that exhibited electron-dense cores when we supplemented the growth media with ferrous iron (**Fig. 5b**), and were distributed as monodisperse spheres throughout the cytosol (**Supplementary Figure 5 c,d**). The electron density maps showed a high similarity to the structure published from encapsulin shells from *M. xanthus* expressed in *E. coli* (pdb 4PT2; EMDataBank EMD-5917^28^ **Figure 5c**). The cutaway views from the encapsulins (blue) furthermore show electron densities associated with docked cargo proteins and most likely biomineralized iron as compared with the inner surface of the shell from *E. coli* (gray) that was mapped in the absence of any cargo (**Fig. 5c** lower row). These data demonstrate that the spherical shape and high non-toxic expression levels make encapsulin very attractive as fully genetically expressed markers for electron microscopy (EM).

**Figure 5:**
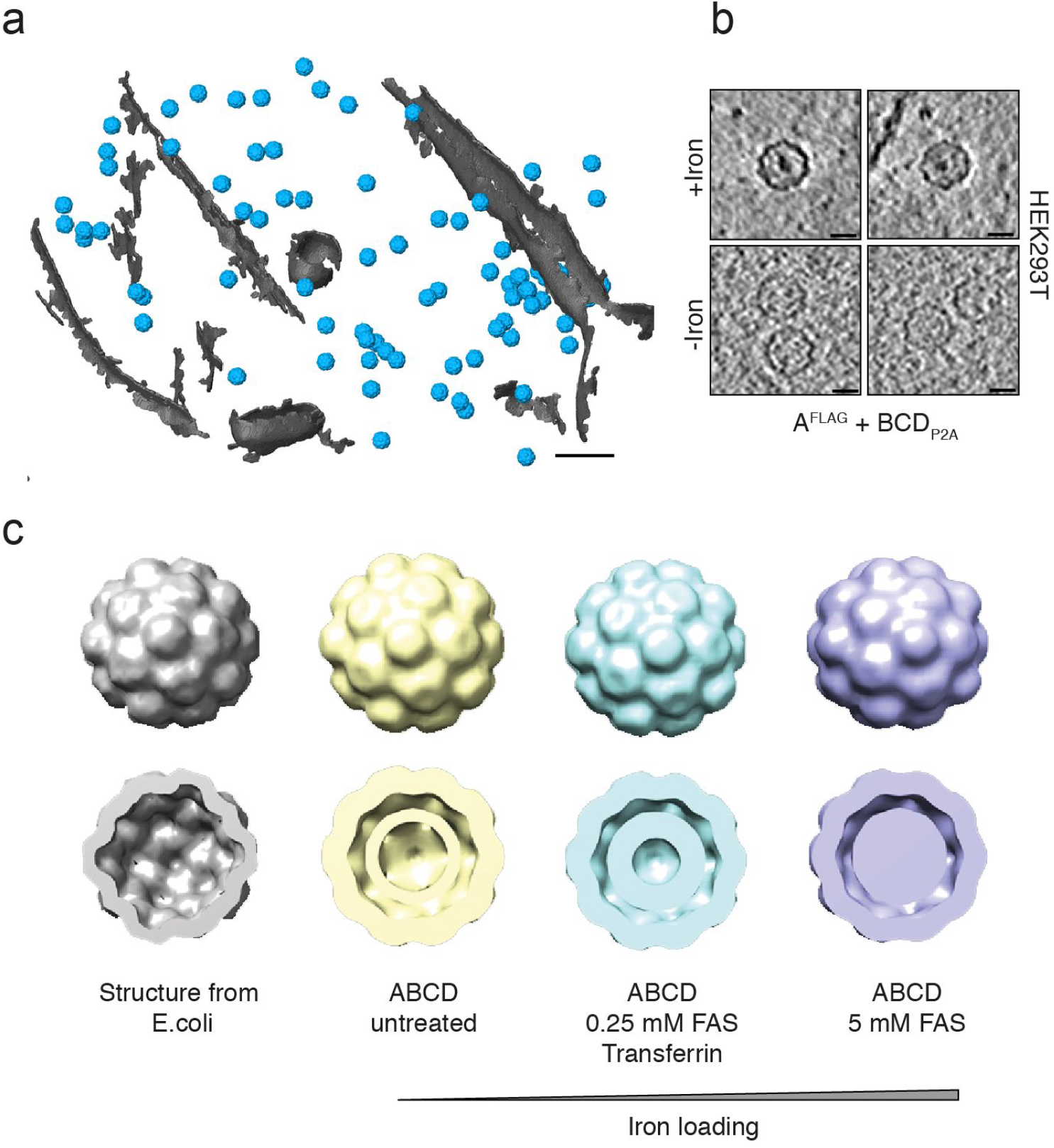
Encapsulins as genetically encoded markers for cryo-electron tomography (Cryo-ET) **(a)** Cryo-ET data from HEK293T cells stably expressing encapsulins together with native ferritin-like cargo proteins (using the dual promoter construct A^FLAG^;BCD^P2A^ shown in Fig. 4a). 3D rendering showing encapsulins in blue and membranes in gray colors. Scale bar:100 nm. **(b)** Example slices from tomograms show encapsulins with and without electron-dense cores from iron-accumulation (treatment with 5 mM FAS for 48 h prior to cryofixation). Scale bars:20 nm. **(c)** Structures (displayed as 2x binned) derived from cryoelectron tomography of nanocompartments assembled in HEK293T cells without (beige), with 0.25 mM FAS and 1 mg/ml human transferrin (cyan) and 5 mM FAS (purple) as compared with the published structure shown in gray (pdb 4PT2; EMDataBank EMD-5917) obtained from *M. xanthus* EncA expressed in *E. coli*^14^. The cutaway views of the encapsulins show electron densities indicating the presence of cargo proteins (beige) and additional iron-deposition (cyan and purple) as compared to published data from the EncA shell^14^ that was obtained in the absence of cargo proteins.

## DISCUSSION

In summary, we genetically controlled multifunctional orthogonal compartments in mammalian cells via expressing N- or C-terminally modified encapsulins, which we found to auto-assemble into abundant nanocompartments which readily encapsulated sets of natural and engineered cargo proteins and enabled size-controlled metal biomineralization.

The efficiency of self-targeting and auto-packaging of the various cargo proteins in mammalian cells was remarkable given that the number of possible protein interactions is even a few-fold higher than in the original prokaryotic host organism^53,54^.

In addition, iron storage inside the capsule via the ferritin-like enzymes B or C targeted to the encapsulins was very efficient, indicating that there was sufficient access to ferrous iron. In contrast, no iron loading of encapsulins heterologously expressed in *E. coli* has so far been shown. We also found that just co-expression of B (or C) with A is sufficient for robust iron storage such that a single-piece reporter construct of just ~2.1 kb in size can be used.

In the context of optimizing T_2_-contrast in MRI, it would certainly be valuable to explore modifications of the outer surface that may control the agglomeration state and thus could modulate the apparent relaxivity of encapsulin ensembles^55^. In this context, it would also be desirable to explore capsid architectures with more storage capacity such as ones with T=7 quasisymmetry known from bacteriophage HK97^28,56^. Furthermore, modifications of the inner surface of the shell may be engineered and/or additional cargo could be designed that could facilitate the nucleation process to support higher iron packing densities or alter environmental parameters (*e.g*. pH and redox potential) to potentially even generate superparamagnetic iron-oxides which possess a substantially larger magnetization^57^. Iterative optimization schemes such as directed evolution could also be employed based on rescue assays from excess iron or magnetic microfluidic sorting and could also be complemented by parallel screens in prokaryotes if enough iron-influx can be achieved there.

Their dense monodisperse distribution, spherical shape and sufficient size, also render encapsulins genetically expressed EM markers in mammalian cells (**Supplementary Video 1**) that are much more readily detectable than ferritins that have been visualized by EM in *E.coli* and yeast^58,59^. In addition, iron-based contrast in encapsulins has the advantage over semi-genetic methods such as metallothionein (MT), miniSOG, erHRP, or APEX/APEX2 that no fixation and delivery of artificial substrates and precipitation of electron-dense material is necessary which may alter cellular structures^42,60–63^. Instead, if iron-based EM contrast is desired, cells expressing the iron-accumulating encapsulins can be grown in regular growth media containing sufficient iron for transferrin-mediated uptake before direct plunge freezing and cryo-electron microscopy (cryo-EM).

For future applications as EM gene reporters in, *e.g*., connectomics research, it would be desirable to generate further encapsulin variants with surface-presented targeting moieties to control their subcellular localization. In this regard, it is of note that - virtue of the self-assembling mechanism - the size of A is just 0.9 kb and that of B just 0.5 kb such that a combined construct is small enough to be carried by viruses optimized for trans-synaptic tracing^64^. It should furthermore be feasible to perform selective detection of encapsulins loaded with split photoactivatable fluorescent proteins via photoactivated localization microscopy (PALM) and combine this with cryo-electron tomography (CET) as was demonstrated for photoactivatable GFP (cryo-PALM)^65^.

Besides allowing influx of metals for size-controlled biomineralizations enabling the applications discussed above, the pore size of ~5 Å inside the encapsulin shell also affords selective passage of small substrates, whereas reaction products may be trapped inside the nanoshell. We have exploited this feature by encapsulating tyrosinase for confined enzymatic production of the toxic polymer melanin and utilized the engineered ‘nanomelanosomes’ as genetically encoded reporters for optoacoustic imaging.

In future applications, encapsulins could thus be used as a versatile reaction chambers for, *e.g*., metabolic engineering of orthogonal reactions in eukaryotic cells. The toolbox for genetically controlled compartmentalization that we introduce here could for instance enable multi-step enzymatic production involving labile and/or toxic intermediates but yielding end-products that may have beneficial intracellular effects or serve as molecular signals upon ‘quantal’ release from the nanocompartment. The approach could for instance also endow genetically modified mammalian cells used for cellular therapies with additional metabolic pathways that may augment their therapeutic efficacy. Complementarily, endogenously produced toxic products could be contained and detoxified in engineered compartments for causal studies or potentially for gene therapeutic approaches.

In addition to the encapsulins presented here, heterologous expression of compartments with different sizes and shapes seem possible, which could offer different sets of endogenous and engineered cargo molecules with different subcellular targeting. These alternative systems would ideally also be orthogonal to each other such that multiplexing (maybe even nesting) of several engineered multicomponent processes could be achieved.

More generally, genetically controlled compartmentalization of multi-component processes in eukaryotic cells - as demonstrated for encapsulins here - is a fundamental biotechnological capability that has profound implications for mammalian cell engineering and emerging cellular therapies.

## Acknowledgements

We are grateful for support from the European Research Council under grant agreements ERC-StG: 311552 (F.S., A.S., H.R., G.G.W.)

## Author contributions

F.S. co-designed the study, generated all constructs, conducted all cell and biochemical experiments, analyzed data and generated figures; C.M. made important contributions to the iron-loading experiments and performed *in vitro* MRI experiments; P.E. conducted and analyzed cryo-EM experiments; A.S. supported cell experiments; H.R. supported animal experiments; M.D., S.B conducted *in vivo* MRI experiments supervised by A.J., J.P. supervised cryo-EM experiments; H.F. and M.H. supervised *in vitro* MRI experiments, V.N. supervised the MSOT experiments. G.G.W. conceptualized and co-designed the study, analyzed data and generated figures, supervised the project, and wrote the manuscript.

## METHODS

### Genetic constructs

Mammalian codon-optimized MxEncA (UniProt:MXAN_3556) MxEncB, MxEncC and MxEncD (UniProt: MXAN_3557, MXAN_4464, MXAN_2410) were custom synthesized by Integrated DNA Technologies and cloned into pcDNA 3.1 (+) Zeocin (Invitrogen) using restriction cloning or Gibson assembly. The MxEncA surface tags (FLAG, StrepTagII or TwinStrepTag) were C-terminally appended using Q5^®^ Site-Directed Mutagenesis (New England Biolabs). N-terminal Myc epitopes were added accordingly to the cargo proteins. Multigene expression of B, C and D was achieved by generating a single reading frame containing all three genes separated by P2A peptides yielding BCD_P2A_. A ‘scarless’ bicistronic construct encoding ^Myc^C and A^STII^ was custom-synthesized by inserting a *Ssp* DnaE mini-intein variant engineered for hyper-N-terminal autocleavage followed by a P2A peptide in between the genes as previously described^66^. For generating stable clones expressing MxEncABCD, MxEncA^FLAG^ was cloned into the Cytomegalovirus promoter (CMV) driven expression cassette of pBudCE4.1 (Invitrogen) and BCD_P2A_ was cloned into the elongation factor 1 alpha promoter (EF1a) driven expression cassette of the vector via restriction cloning. To generate Adeno-associated viruses (AAV) enabling multigene expression of MxEncA^FLAG^ and MxB-Mms7ct, two strategies were employed: MxEncA^FLAG^ was cloned upstream of a ECMV internal ribosome entry site (IRES) whereas MxB-Mms7ct was inserted downstream. The second approach employs MxB-Mms7ct followed by a P2A peptide and MxEncA^FLAG^. The two cassettes were subcloned into pAAV-CamKIIa (https://www.addgene.org/26969/) with BamHI and EcoRI. AAVs were custom prepared by the UNC Vector Core of the University of North Carolina at Chapel Hill. To test the bicistronic expression constructs used for the AAVs in HEK293T cells, the cassettes were also sub-cloned into the pcDNA 3.1 (+) Zeocin with EcoRI and NotI. To target PAmCherry1 and mEos4b as cargo to the encapsulin nanocompartments, the fluorescent proteins were C-terminally fused to 2 x GGGGS linkers followed by the minimal encapsulation signal LTVGSLRR (EncTag). For complementation of split PAmCherry1 inside the encapsulin nanoshell, amino acids 1-159 of PAmCherry1 were fused to MxEncC via a 2 x GGGGS linker and amino acids 160-236 of PAmCherry1 were directly fused to the C-terminus of MxEncB. For complementation of a split luciferase, the split part LgBit (NanoBiT system, Promega) was fused C-terminally to MxEncC via a 2 x GGGGS linker. SmBit was directly fused to the C-terminus of MxEncB. SmCSE^44^ (UniProt: Smal_0489) and APEX2^42^ were fused to 2 x GGGGS linker followed by the minimal encapsulation signal. Mammalian codon-optimized *Bacillus megaterium* tyrosinase (BmTyr) was C-terminally appended to ^Myc^D separated by 2 x GGGGS linker in custom gene synthesis. C-terminally FLAG-tagged *Mus musculus* Zip14 was inserted into pcDNA 3.1 (+) or pIRES2-ZsGreen1 via restriction cloning. To yield secreted encapsulins, MxEncA^STII^ was N-terminally fused to a BM40 mammalian secretion peptide. In order to generate enacpsulin derivatives featuring C-terminal acidic peptides of magnetotactic bacteria Mms proteins that are implicated in mediation of magnetite formation either the C-terminal peptide of Mms6 (YAYMKSRDIESAQSDEEVELRDALA) or Mms7 (YVWARRRHGTPDLSDDALLAAAGEE) of *Magnetospirillum magneticum* were fused either to the inward-facing N-terminus of MxEncA^FLAG^ or to the C-terminus of either the MxEncB or C using Q5^®^ Site-Directed Mutagenesis. For a complete list of the genetic constructs featuring their composition refer to **Supplementary Table 1**.

### Cell culture, transfection and generation of stable cell lines

Low passage number HEK293T (ECACC: 12022001) and CHO cells were cultured in advanced DMEM with 10 % FBS and Penicillin-Streptomycin at 100 μg/ml at 37 °C and 5 % CO_2_. Cells were transfected with X-tremeGENE HP (Roche) according to the protocol of the manufacturer. DNA amounts (ratio shell to cargos) were kept constant in all transient experiments to yield reproducible DNA-Lipoplex formation. To generate a stable HEK293T cell line expressing MxEncABCD, cells were transfected with pBudCE4.1 MxEncABCD and stable transfectants were selected with 300 μg/ml Zeocin (InvivoGen).

### Protein expression and lysis

Cells were harvested between 24 and 48 h post-transfection. Cells were lysed with M-PER Mammalian Protein Extraction Reagent (Pierce Biotechnology) containing a mammalian protease inhibitor cocktail (SIGMA P8340, Sigma-Aldrich) according to the protocol of the manufacturer in all experiments using FLAG-tagged encapsulins. For lysis of cells expressing StrepTagII-modified encapsulins, cells were resuspended in buffer W (150 mM NaCl, 100 mM Tris-Cl, pH 8.0) and exposed to 4 freeze-thaw-cycles in LN_2_. After spinning down cell debris at 10,000 x g for 15 min, cell lysates were kept at 4 °C for downstream analyses. Protein concentrations of lysates were determined measuring OD at 280 nm.

### Co-Immunoprecipitation of Encapsulins

Cell lysates were incubated with Anti-FLAG® M2 Magnetic Beads or Anti-FLAG® M2 affinity gel (SIGMA M8823 and A2220, Sigma-Aldrich) according to the protocol of the manufacturer. After binding, the magnetic beads were washed 4 times on a magnetic separator rack (DYNAL separator, Invitrogen) with M-PER buffer. Bound FLAG-tagged encapsulins were eluted using M-PER buffer containing 100 μg/ml FLAG-peptide (SIGMA F3290, Sigma-Aldrich). In the case of enacpsulins with an external StrepTagII, MagStrep “type3” XT beads or Strep-Tactin®XT resin (IBA Lifesciences) was used according to the protocol of the manufacturer. Proteins were eluted using Buffer BXT containg 50 mM Biotin. To analyze the eluted proteins, samples were mixed with SDS-PAGE sample buffer and incubated at 95 °C for 5 min. Samples were loaded onto pre-cast 12 % Bio-Rad Mini-PROTEAN® TGX™ (Bio-Rad Laboratories) gels and run for 45 min at 200 V. Accordingly, gels were either directly silver-stained using SilverQuest™ Silver Staining Kit (Novex) according to protocol of the manufacturer or immuno-blotted onto PVDF membranes. After blotting, membranes were blocked in 5 % non-fat milk in TBS for 1 h at room temperature. Subsequently, membranes were incubated in TBS containing 5 % non-fat milk and 1 μg/ml Monoclonal ANTI-FLAG^®^ M2 antibody (SIGMA F1804, Sigma-Aldrich) or 1 μg/ml Anti-Myc Tag Antibody clone 9E10 (05-419, EMD Millipore) for 2 h at room temperature. After five washing cycles with TBS, membranes were incubated with anti-mouse IgG HRP-conjugate (SIGMA A5278, Sigma-Aldrich) for 1 h at room temperature in 5 % non-fat milk in TBS. Protein bands were detected using Amersham ECL Prime Western Blotting Detection Reagent (GE Healthcare Bio-Sciences AB) on a Fusion FX7 / SL advance imaging system (Peqlab Biotechnologie GmbH).

### Blue Native gel electrophoresis and on-gel analyses

For detection of native encapsulin nanocomparments, the NativePAGE™ Novex ® Bis-Tris Gel System (Novex) was used. Either eluted material from the Co-IP/purification or whole cell lysates of cells expressing encapsulins in NativePAGE™ Novex® sample buffer were loaded onto pre-cast NativePAGE™ Novex® 3–12% Bis-Tris gels. The total protein amount of whole cell lysates loaded per well was adjusted to ~1–3 μg. Blue native (BN) gels were run for 90 – 180 min at 150 V according to the protocol of the manufacturer. Gels loaded with samples from Co-IP/purification were silver-stained using SilverQuest™ Silver Staining Kit (Novex) or Coomassie-stained using Bio-Safe™ Coomassie Stain (Bio-Rad Laboratories). For protein detection, gels loaded with whole-cell lysate samples were Coomassie-stained accordingly. For detection of iron-containing proteins, gels loaded with samples containing iron loaded encapsulins were Prussian Blue (PB) stained. Briefly, gels were incubated in 2 % potassium hexacyanoferrate(II) in 10 % HCl for 45 min. For 3,3’-Diaminobenzidine (DAB)-enhancement (DAP PB), gels were washed three times with ddH_2_O and incubated in 0.1 M phosphate buffer pH 7.4 containing 0.025 % DAB and 0.005 % H_2_O_2_ until dark-brown bands appeared. To stop DAB polymerization, gels were washed three times with ddH_2_O. For detection of fluorescent signals from native encapsulin bands (fluorescent cargos: mEos4b, PAmCherry1, split PAmCherry1 or mineralized CdS), unstained BN gels were imaged on a Fusion FX7 / SL advance imaging system (Peqlab Biotechnologie GmbH) using the UV fluorescence mode. For on-gel detection of luminescence signal generated by encapsulated split NanoLuciferase, unstained BN gels were soaked in 1 ml of Nano-Glo® Luciferase substrate (Nano-Glo® Luciferase Assay, Promega) and imaged on a Fusion FX7 / SL advance imaging system (Peqlab Biotechnologie GmbH) in chemiluminescence mode. For whole-cell lysate luminescence detection, cell lysates were mixed with the substrate at a 1:1 ratio and luminescence readings were taken on a Centro LB 960 (Berthold Technologies) at 0.1 s acquisition time. For detection of APEX2 peroxidase activity inside encapsulins, unstained BN gels were incubated in 0.1 M phosphate buffer pH 7.4 containing 0.025 % DAB and 0.005 % H_2_O_2_ for 15 min until black bands appeared on the gel. For microscopic detection of DAB polymerization in cells expressing APEX2-loaded encapsulins, cells were fixed in 4 % PFA in PBS for 15 min. Subsequently, cells were incubated in 0.1 M phosphate buffer pH 7.4 containing 0.025 % DAB and 0.005 % H_2_O_2_ for 5 min. The reaction was stopped by washing three times with PBS. For the on-gel detection of melanin generation inside encapsulins, gels loaded with whole cell lysates of HEK293T cells expressing encapsulins loaded with tyrosinase were incubated in PBS containing 2 mM L-tyrosinase and 100 μM CuCl_2_ for 1 h at 37 °C until a black encapsulin band became visible.

### Microscopy of complementation of split PAmCherry1 inside encapsulins

Cells transfected with C-PAs1 and B-PAs2 with or without A^FLAG^ were seeded onto 8-well Poly-L-Lysine-coated microscopy chips (Ibidi). 36 h post transfection, live cell confocal microscopy was conducted on a Leica SP5 system (Leica Microsystems). For photoactivation of split PAmCherry1, samples were illuminated with a 405 nm laser for 60 s at 40 % laser power. The signal of complemented split PAmCherry1 was excited using the 561 nm laser. To quantify the complementation of split PAmCherry 1 with or without the encapsulin shell, the ratio of the total mean fluorescence after photoactivation divided by the signal before was calculated. ImageJ was used to quantify mean fluorescence values from randomly chosen areas on the well.

### Multispectral Optoacoustic Tomography (MSOT)

Optoacoustic images of cells co-expressing A^STII^ and ^Myc^D-BmTyr were acquired on an inVision 256-TF system (iThera Medical GmbH). Briefly, ~10^7^ HEK293T cells co-expressing the genes treated with 10 μM CuCl_2_ and 1 mM L-tyrosine 24 h prior to the measurement were detached using trypsin, washed with PBS embedded into 1 % low melting agar yielding a tubular phantom of ~ 300 μl volume. The cell phantom and an ink phantom (OD=0.2) were placed in a custom-built sample holder and optoacoustic images were acquired for the range of wavelengths between 690 and 900 nm. Signals were reconstructed using ViewMSOT software suite (iThera Medical GmbH) and linearly unmixed using a reference spectrum for melanin.

### Magnetic Sorting

Cells were washed twice with PBS, detached with accutase (Sigma-Aldrich) and resuspended in DPBS supplemented with 10% fetal bovine serum (Gibco) prior to sorting. For magnetic sorting, columns filled with ferromagnetic spheres (MS columns, Miltenyi Biotec) were placed in an external magnetic field (OctoMACS separator, Miltenyi Biotec) and equilibrated with 1 ml DPBS containing 10% FBS. The column was loaded with cells and washed with 0.5 ml DPBS; the flow-through was collected as one fraction. After removing the column from the magnetic separator, cells were eluted with 1 ml DPBS. The total number of cells before sorting as well as the cell numbers in flow-through and eluate were determined with a Countess II FL Automated Cell Counter (Life Technologies).

### MRI of cells

MR images were acquired at a Bruker BioSpec 94/20USR, 9.4 T system equipped with a RF RES 400 1H 112/072 Quad TR AD resonator. For T_2_* measurements of cell pellets, 4*10^6^ HEK293T cells were seeded 24 h prior to transfection on Poly-L-lysine-coated 10 cm cell culture dishes. 24 h post transfection, ferrous ammonium sulfate (FAS) was added to the medium yielding a concentration of 1 mM. 24 h post iron addition, cells were washed three times with DPBS and detached with Accutase and centrifuged at 500 x g for 4 min. The pellets were resuspended in 800 μL DPBS and transferred to cryobank vials (Thermo Scientific Nunc) containing 50 μl of solidified 1% agarose at the bottom. Cells were then spun down at 2,000 x g for 2 min and immediately used for MRI. T_2_* measurements were conducted in a custom-made holder filled with DPBS to avoid susceptibility artefacts. T_2_*-values were calculated based on a multiple gradient echo (MGE) sequence with a TR of 800 ms, 12 echoes with an echo spacing of 4.5 ms (3.5 – 58.5 ms), a flip angle of 50°, field of view of 65 x 65 mm and a matrix size of 256 × 256. Relaxation rates were calculated with the *Image Sequence Analysis Tool* from Bruker BioSpin MRI GmbH.

### *In vivo* MRI

HEK293T cells (~4*10^6^) were seeded onto Poly-L-Lysine-coated 10 cm cell culture dishes. 24 h after seeding, cells were transiently transfected at 70–80 % confluency with DNA constructs encoding either A^FLAG^+B^M7^ or A^FLAG^+mEos4b-EncTag, as well as for both conditions Zip14 at 5 % of the total DNA amount using X-tremeGENE™ (Roche). 24 h post transfection, the cell culture medium was replenished with fresh medium containing 1 mM FAS. 24 h after incubation with FAS, cells were washed gently 3 times with PBS, detached from the culture dishes after 5 minutes of treatment with a 1:1 solution of Accutase (Sigma) and Trypsin, centrifuged for 5 min at 1,200 x g and resuspended in growth media. Cell suspensions were backfilled into two injection cannulae (28 gauge, Plastics One, Roanoke, VA, USA) connected via plastic tubing to 25 μl Hamilton glass syringes clamped in a remote dual syringe pump (PHD 22/2,000; Harvard Apparatus, Holliston, MA, USA). Injection cannulae were then lowered into bilateral guide cannulae (22 gauge, Plastics One, Roanoke, VA) that were previously implanted in Sprague-Dawley rats^67^. Rats were then centered in the bore of a 7T 20 cm inner diameter, horizontal bore magnet (Bruker BioSpin MRI GmbH, Ettlingen, Germany) and gradient echo scans (FOV = 2.5 cm × 2.5 cm, matrix size = 256 × 256; 7 slices with 1 mm slice thickness) were taken at a TR = 800 ms and different echo times (5, 10, 15, 20, 25 ms) to compute relaxation rate maps and perform ROI analysis (circular ROIs of 1 mm diameter placed on injection sites) using custom routines in Matlab (Mathworks, Natick, MA, USA).

### Cell viability assays

Iron-related cytotoxicity was monitored via the Roche Cytotoxicity Detection Kit (LDH) (Roche Diagnostics) according to the protocol of the manufacturer. Briefly, 7,5*10^5^ HEK293T cells were seeded on Poly-L-lysine-coated 24-well plates. 24 h post seeding, cells were transfected with different combinations of genes using X-tremeGENE HP (Roche). The Zip14 DNA amount was kept constant in all samples expressing Zip14 (5 % of total DNA). For expression of combinations of A^FLAG^ with cargo proteins, 60 % of the total DNA amount was encoding A^FLAG^ and the remaining 35 % were used for the respective cargo molecule. 24 h post transfection, FAS was added to the medium from a 100 mM stock solution yielding a final concentration of 2.5 mM. 24 h post addition of FAS, cells were assayed for LDH release. In order to evaluate gene-related toxicity in the absence of iron, the assay was performed accordingly but without iron supplementation and cells were assayed 48 h post-transfection. The Luciferase-based viability assay (RealTime-Glo™ MT Cell Viability Assay, Promega) was performed according to the protocol of the manufacturer in 96-well plate format as an end point measurement. Luminescence readings were taken on a Centro LB 960 (Berthold Technologies) at 0.5 s acquisition time.

### Electron Microscopy

#### Sample Preparation and Cryo FIB Milling

Purified protein samples were applied on glow-discharged R1.2/1.3 grids (Quantifoil), plunge-frozen using a Vitrobot Mark IV (FEI, settings: blotforce = 20, blottime = 5 s) and stored under liquid nitrogen (LN_2_) until use. HEK293T cells stably expressing MxEncABCD were grown in DMEM + 10% FBS and GlutaMAX in the presence of penicillin-streptomycin (P/S) and Zeozin (‘growth media’). 0.25 mM FAS in the presence of transferrin (2 mg/mL) or 5 mM of FAS were used to increase the fraction of encapsulin compartments with electron dense cores over those expressed in cells grown without iron supplementation. For each of the conditions, cells were dissociated using 2.5% trypsin and counted. 4 uL of a suspension containing 150 cells/μL were plated on R1/4 carbon or SiO_2_ grids (Quantifoil), which had been coated with an additional layer of carbon (20 nm), glow-discharged and coated with PDL. After 24 h, the medium was changed to growth media + 10% glycerol and the cells were incubated at 37 °C for 10 min. Cells were vitrified using a Vitrobot Mark IV (FEI, settings: blotforce = 12, blottime = 10 s, draintime = 1 s) and stored under LN_2_ for further processing. Grids were clipped in custom-made AutoGrids and ultrathin lamellae of randomly chosen cells were cut according to published procedures^68^ using a focused ion beam microscope (FEI, model: DB Quanta 3D FEG) equipped with a cryo-system (Quorum Technologies, model: PP3000T) and a custom-made cryo stage (MPI for Biochemistry, Martinsried).

#### Transmission Electron Microscopy

Single particle grids were imaged at a nominal defocus of −5 μm and an EFTEM magnification of 42000x on a transmission electron microscope (FEI, model: Titan Krios, FEG 300 kV) with a post-column energy-filter (Gatan, model: 968 Quantum K2). Images were recorded with a direct detection camera (Gatan, model: K2 Summit) using the SerialEM software package^69^. Tomography acquisition on cut lamellae was performed at a defocus of −5 μm. The tilt range depended on the pre-tilt and was typically −50° to +70° with a tilt increment of 2°. The total dose was kept below 90 e–/Å2 and the unbinned pixel size was 3.42 Å. Individual frames were aligned using MotionCorr2^70^. Tomograms were reconstructed with IMOD^71^ and processed using the PyTom software package^72^ as well as Matlab (MathWorks).

#### Template Matching, Subtomogram Averaging and Visualization

For each of the three iron-loading conditions (0 mM FAS, 0.25 mM FAS + transferrin and 5mM FAS) the encapsulin nanospheres as well as ribosomes (exemplary tomogram Fig. 5a) were localized by template matching on 4 times binned tomographic data using down-filtered templates constructed from manually picked (~200) particles. Subvolumes of unbinned particles were reconstructed, CTF-corrected and averaged in RELION applying icosahedral symmetry (I3) and using the average of the particles as an initial reference with a spherical mask. Averaged volumes were filtered to their respective resolution and rendered with UCSF Chimera (http://www.cgl.ucsf.edu/chimera) or Amira (FEI).

---

**TOC:**
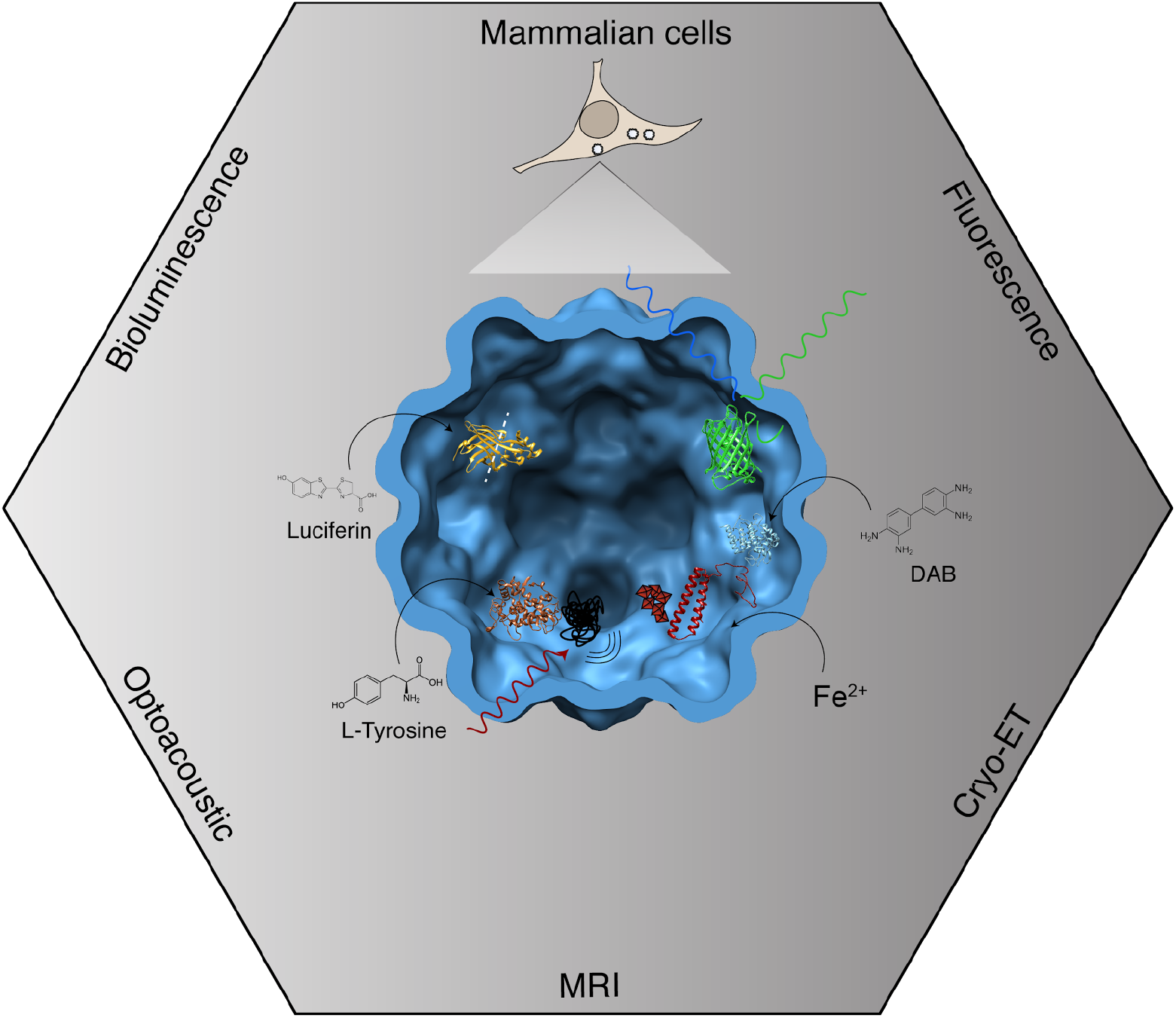
Genetically controlled compartmentalization of mammalian cells via eukaryotically expressed encapsulins enables sequestration of multi-component processes such as enzymatic reactions and detoxifying metal biomineralization for applications in multimodal imaging and mammalian cell engineering.

